# A Novel Deep Learning Method using Contrastive Learning Enables Interpretable Outputs from Relationships Between Gene Expression and Histone Modification

**DOI:** 10.1101/2025.09.04.674139

**Authors:** Abdul Munif, Amitava Datta, Zhaoyu Li, Max Ward

**Affiliations:** School of Physics, Mathematics, and Computer Sciences, The University of Western Australia, Australia; School of Biomedical Sciences, The University of Western Australia, Australia

**Keywords:** Epigenetics, Histone Modifications, Gene Expression, Deep Learning, Pairwise Ranking, Neural Networks, Machine Learning

## Abstract

Current methods for predicting gene expression from histone modifications rely on arbitrary binary classification thresholds such as the median to distinguish between high and low expression for genes. This approach lacks biological justification, creates dataset-dependent classifications, and ignores the relative regulatory relationships between genes that are often more biologically meaningful than absolute cutoffs.

We introduce a novel pairwise ranking approach that compares relative expression levels between gene pairs based on their histone modification patterns, eliminating arbitrary threshold selection. Evaluation using the REMC E066 and GSE76344 datasets showed that our pairwise classification framework demonstrated consistent ranking performance in both datasets. The ablation study found that only a subset of histone modification types is necessary, giving evidence of redundancy in the biological code.

## Introduction

Epigenetics is a field that explores heritable changes in gene expression without alterations to the DNA sequence. It is increasingly significant due to its implications in various diseases, cancer, cognitive dysfunction, and autoimmune disorders (1). These changes are influenced by environmental factors such as exposure to heavy metals, pesticides, and tobacco smoke, as well as nutritional elements, with the potential to persist across multiple generations. Epigenetic modifications occur through mechanisms like DNA methylation, chromatin modification, imprinting, and the involvement of non-coding RNAs, each playing a critical role in gene regulation and disease development (2, 3).

Recent advancements have enhanced our understanding of the complex molecular landscape of diseases like cancer. However, traditional methods often fall short in capturing the nonlinear relationships present in biological data. This limitation has led to the exploration of deep learning models, which offer the potential to uncover more complex patterns and correlations between histone modifications and gene expression. Models like DeepChrome (4) and its successor, DeepChrome 2.0 (5), utilize convolutional neural networks and generative adversarial networks (GANs) to improve prediction accuracy and provide deeper insights into the epigenetic regulation of genes.

The research proposes a novel approach to gene expression prediction, framing it as a pairwise ranking problem, where the relative expression levels of two genes are compared based on their histone marks. Based on learning-to-rank approaches (6–8), we aim to develop a neural network architecture for pairwise ranking. The primary objective is to identify which combinations of histone marks most strongly predict relative expression differences between gene pairs.

Deep learning is famous for its capacity to find complex relationships between the data. Singh et al. (4) gave a novel computational framework for predicting gene expression based on histone modification data. The framework, called DeepChrome, was built based on convolutional neural networks (CNNs). While DeepChrome showed competitive performance with traditional methods like Support Vector Machines and Random Forests, subsequent studies like ShallowChrome (9) have suggested that the performance gains of deep learning approaches in this domain may be more modest than initially reported.

Singh et al. (10) later developed an attention-based approach called AttentiveChrome. Attention was proposed to help AttentiveChrome identify relevant positions within chromatin markings. It consists of five modules which are Bin-level LSTM encoder for each histone modification (HM) mark, Bin-level Attention on each HM mark, HM-level LSTM encoder that encodes all HM marks, HM-level Attention over all the HM marks, and the classification module. The drawbacks of AttentiveChrome are that the results rely on correlation with only H3K27ac for validation, which is available for just 3 out of 56 cell types. The model also assumes a sequential ordering of histone marks even though no clear biological ordering exists.

Chen et al. (11) presents neural network layers with selfattention mechanism and transfer learning, called TransferChrome, to improve prediction accuracy, especially in across cell lines. The transformer architecture was used as self-attention mechanism for aggregating global information. While TransferChrome achieved marginally higher AUC scores than existing methods (0.043 improvement over DeepChrome, 0.023 over AttentiveChrome, and only 0.006 over ConvChrome), these gains are modest and may not justify the added architectural complexity. The arbitrary selection of E085 as the source domain for all cross-cell line experiments lacks justification and undermines the generalizability claims of the transfer learning approach.

Frasca et al. (9) introduce ShallowChrome, a novel computational pipeline for the accurate and interpretable prediction of gene expression based on histone modifications. It uses a feature extraction process based on peak calling from ChIP-Seq data. This method identifies significant genomic regions where histone marks are enriched and uses these features for gene expression prediction in binary classification. Because ShallowChrome models use a logistic regression model, it can be interpreted straightforwardly from its model parameters. While this linear regression coefficient claimed to provide meaningful biological interpretability, these weights merely sho statistical association rather than causal mechanism. The validation using only one gene (PAX5) across three cell types may not provide sufficient evidence about extracting regulatory insights across thousands of genes and dozens of epigenomes.

The main challenge encountered by the deep learning approach is interpretability (4, 9–11). Deep learning is often seen as a black box model, making it hard to interpret the learned interactions and the effect of histone modifications on gene predictions. The inability to achieve accurate predictions across different cell lines has also been studied (11). The usage of binning strategies, where separate models were trained for each bin of DNA regions, could not figure out the interactions between neighboring bins. This is important for understanding the spatial distribution of histone modifications. A more complex model can improve the model accuracy, but also increase the model interpretability. It is difficult to interpret the model’s predictions and understand the contribution of different histone modification marks to gene expression (9, 11).

## Related Work

### Pairwise Ranking

Burges et al. (6) proposed a new approach called RankNet to improve the ranking accuracy and efficiency in search engines. The goal of their approach was to order a set of documents returned for a given query based on their relevance. The main components of the approach are probabilistic cost function based on pairs of examples, the use of neural networks to model the underlying ranking function to capture complex relationships in the data, and modified backpropagation algorithm for training on pairs of examples. This resulting ability to learn complex ranking function and improved performance with increasing training set size and handled ties effectively.

An additional method called LambdaRank and LambdaMART has been proposed to improve the search result quality, optimizing information retrieval measures (e.g. NDCG) (7). The LambdaRank builds upon RankNet and directly specifies gradients (lambdas) instead of deriving them from a cost function. The LambdaMART combines LambdaRank with MART (Multiple Additive Regression Tree). It uses gradient boosting with regression tree. These proposed methods can be easily adapted to optimize various IR (Information Retrieval) measures.

The pairwise learning-to-rank approach called DirectRanker has been proposed by Köppel et al. (8). This approach generalizes the RankNet architecture, making it simpler. The main components of the algorithm lies in a neural network structure with two identical subnets that process document features, followed by taking their difference and passing it through an output neuron. The model is proven to be reflexive, antisymmetric, and transitive, allowing for simplified training and improved performance. These properties enable training on only neighboring relevance classes and avoid the need to explicitly train on document equality within classes.

## Dataset

We use two datasets below to test our proposed approach.

### REMC database

The Roadmap Epigenome Project (REMC) (12) is a public database of human epigenomic data containing histone modifications and gene expression data from mRNA-Seq from 56 cell types of healthy people. In Singh et al. (10), they used all cell types and five core histone modification marks (H3K4me3, H3K4me1, H3K36me3, H3K9me3, H3K27me3). To make comparison with our proposed method, we only use the E066 (adult liver cell). In addition, we use all available histone modification marks.

### GSE76344 dataset

The GSE76344 is a dataset that contains histone modifications cell lines gathered from Zheng et al. (13). This dataset contains multiple histone modification markers such as H3K27ac, H3K27me3, H3K4me3, H3K9ac, and H3K9me3. The histone modifications on REMC dataset and GSE76344 dataset shown in Table 1. There are three common histone marks (H3K4me3, H3K27me3, and H3K9me3).

**Table 1.**
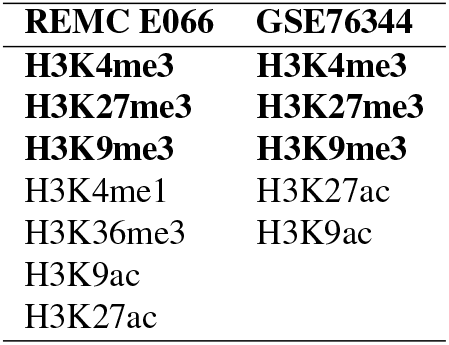
Histone marks found in REMC E066 dataset and GSE76344 dataset

## Proposed Method

We have developed a comparative model to assess the relative gene expression levels between two genes. This model utilize a pairwise ranking approach, evaluation the histone marks of two genes to determine which gene are having a higher level of expression. The model was built similar with DirectRanker (8), which using neural network approach to do pairwise ranking. The pairwise ranking architecture can be seen on Figure 1.

**Fig. 1.**
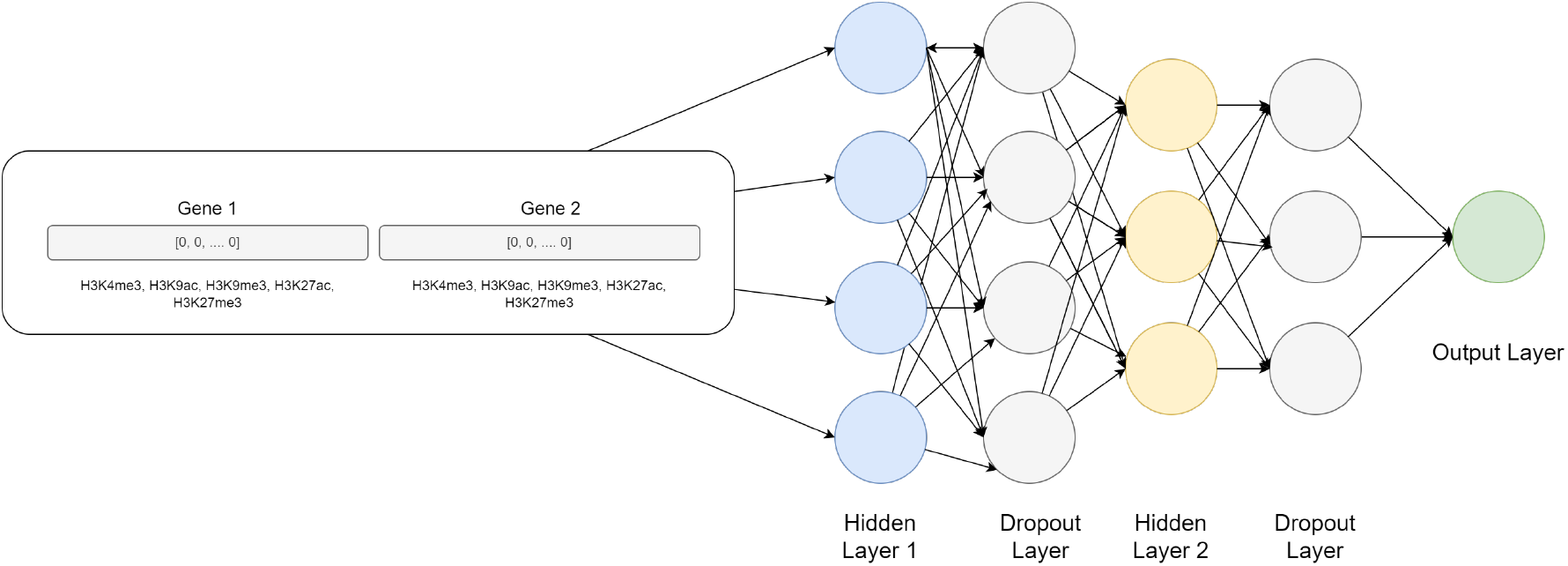
Pairwise Ranking Neural Network Architecture

The pairwise ranking with neural network consists of the following layers:

1. Input layer The model’s input contains the epigenetic profiles of two genes, each characterized by five distinct histone modification markers. We have implemented two feature representation strategies to accommodate varying levels of computational complexity and data dimensionality:
  a. **Full Feature Representation**: In this approach, we utilize the complete set of features, resulting in a high-dimensional input vector. Each gene is represented by 20,000 features, encompassing the full spectrum of histone modification data across the gene body and its proximal regulatory regions.
  b. **Reduced Feature Representation**: To address potential computational constraints and to mitigate the curse of dimensionality, we have also developed a low-dimensional feature representation. In this case, each gene is characterized by a condensed set of 100 features, which are designed to capture the most salient aspects of the histone modification patterns.
  For both representation strategies, the feature vectors of the two genes under comparison are concatenated to form a single input array. Consequently, the model processes either a 40,000-dimensional vector (in the case of full feature representation) or a 200-dimensional vector (for the reduced feature set). This concatenation allows the model to simultaneously consider the epigenetic profiles of both genes, facilitating direct comparison and relative ranking of their expression levels. The input layer will become a single array with size *N*_*k*_, where *k* = {200, 40000}.
2. Hidden layer 1 The next layer is hidden layer with size *k* × *n*_1_, where *n*_1_ is the size of output of first hidden layer. This layer will apply the linear transformation to the incoming data, mapping the input features to a new feature space. The linear transformation can be expressed as:

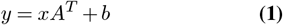

where *y* denotes the output of the first hidden layer, *x* i the weight matrix of *k* × *n*_1_, *A* represents the input featur vector, and *b* is the bias vector.
3. Rectification 1 We apply the non-linearity function called LeakyReLU. This will introduce nonlinearity in neural network and can help with vanishing gradient problem during training. It will enable the neural networks to learn more complex relationships in the data. The LeakyReLU can be expressed as:

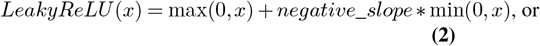

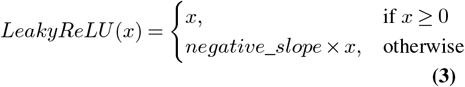
4. Dropout layer 1 The next layer is dropout layer. This layer will randomly zeroes some of the elements of the input tensor with probability *p*. The dropout layer can avoid overfitting in the model, a common challenge in deep learning models. The dropout layer promotes the learning of more robust and generalized representations, and will enhance the model’s performance on unseen data.
5. Hidden layer 2 Subsequent to the dropout layer, the data is processed through a second hidden layer. This layer is characterized by dimension *n*_1_ × *n*_2_, where *n*_1_ is the input size (output of the previous layer) and *n*_2_ represents the number of output units in this second hidden layer.
6. Dropout layer 2 To further enhance the model’s generalization capabilities, an additional dropout layer is incorporated following the second hidden layer. This layer employs the same dropout probability *p* as the first dropout layer, reinforcing the regularization effect throughout the network architecture.
7. Output layer The final layer of the network architecture is a fully connected layer that performs a linear transformation from the input data followed by sigmoid function to do binary classification. The size of fully connected is *n*_2_ × 1, where *n*_2_ is the output size of the previous layer and 1 is the number of output neuron that represent the binary classification. The sigmoid function is used to make class predictions by mapping it to a probability value in the range [0, 1], and can be expressed as:

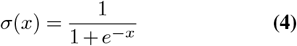

This probability output is then thresholded at 0.5 to make the final binary classification decision. The final result will be interpreted as follows:

- ‘1’ if the first gene expression level is higher than the second gene expression level.
- ‘0’ if otherwise.

We also develop another ranking method by adding the preprocessing layer. This was inspired by DirectRanker approach (8). We added the various feature extraction layer based to extract the meaningful features from histone modification. These are:

- Random Projection
- Transformer
- Auto Encoder (using fully connected layer)
- Auto Encoder (using convolution layer)

The pairwise ranking architecture with additional extraction can be seen on Figure 2. The *n*_1_ and *n*_2_ is a certain feature extraction layer.

**Fig. 2.**
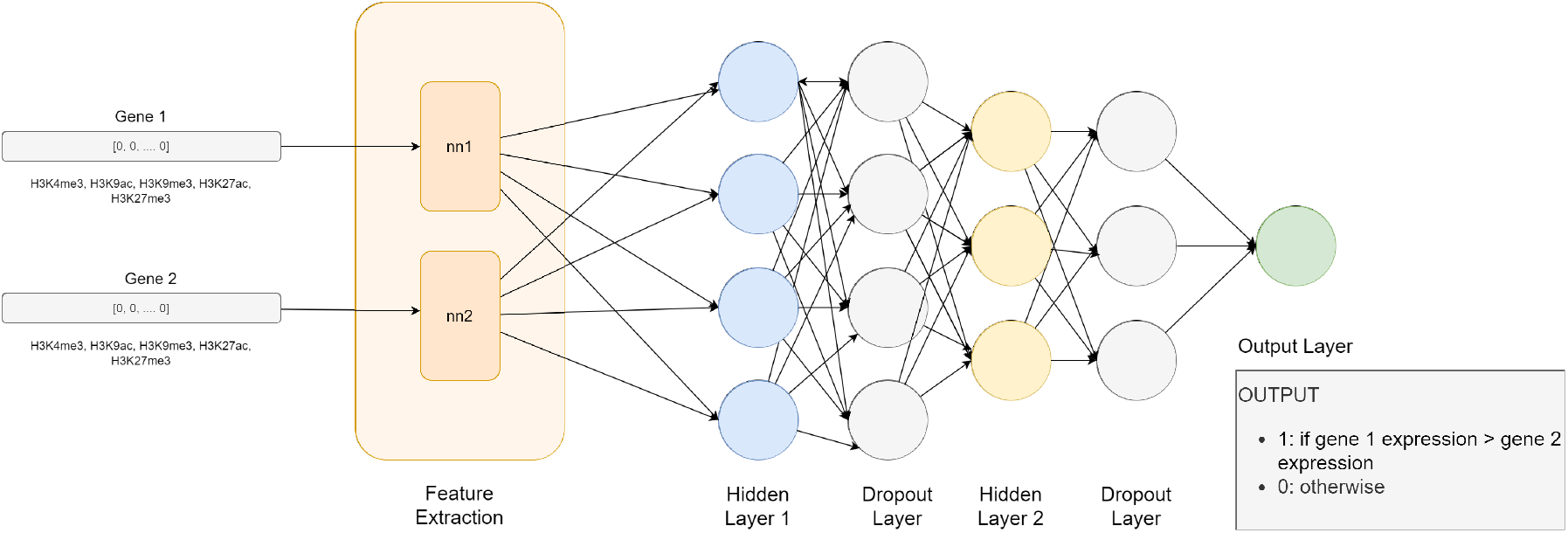
Pairwise Ranking with Feature Extraction Layer

## Experiment Setup

We will explain the data preprocessing step and the experiment setup we are using in this research.

### Data Preprocessing

#### REMC E066 dataset

We follow the data preprocessing or feature extraction of REMC E066 from Singh et al. (4). First, they divided 10,000 basepair (bp) DNA Region (+/-5,000 bp) around the TSS (transcription start site). Then, this region will be binned into bins of length 100 bp and extract the mean of gene expression level in the bins. They consider five histone marks as explained in Table 1. This makes the input for each gene a 5 × 100 matrix, where columns represent different bins and rows represent histone modifications.

#### GSE76344 dataset

In our study, we adapted the feature extraction methodology established by Singh et al. (4), with modifications tailored to the specific characteristics of the GSE76344 dataset. Our approach focuses on a more concentrated genomic region and accommodates the binary nature of the available histone modification data. The overall procedures are explained in Figure 3.

**Fig. 3.**
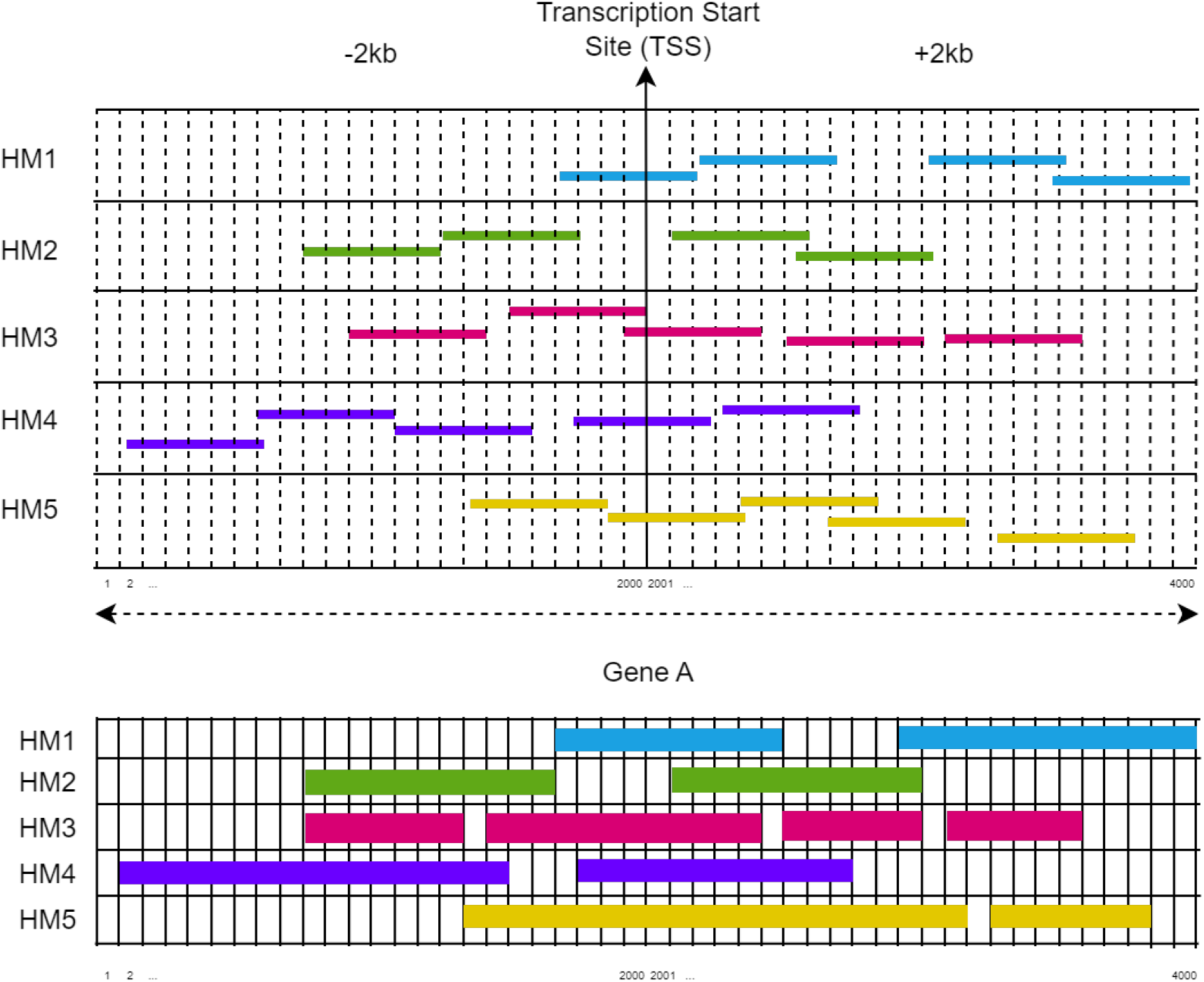
Feature extraction on GSE76344 dataset

1. **Genomic Region of Interest**: We take a 4,000 base pair (bp) region centered on the Transcription Start Site (TSS), extending 2,000 bp upstream and 2,000 bp downstream. This region was selected to capture the most proximal and potentially influential epigenetic features.
2. **Histone Modification Encoding**: Unlike the signal peak data available in Singh et al. (4) study, the GSE76344 dataset provides binary information on histone mark presence. We encoded this information as follows:
  - ‘1’ indicates the presence of a specific histone modification at a given location.
  - ‘0’ indicates the absence of histone modification at a given location
3. **Feature Array Construction**: For each of the five histone modifications under consideration, we generated a 1 × 4000 binary array for each histone mark, resulting in a composite 5 × 4000 matrix for the complete set of histone marks.

The resulting feature matrices are adaptable to different neural network architectures:

1. DeepChrome Architecture: The 5 × 4000 matrix format is directly compatible with the DeepChrome model, preserving the spatial relationships between histone modifications and genomic positions.
2. Pairwise Ranking Architecture: For this architecture, the 5 × 4000 matrix is flattened into a 1 × 20000 vector, facilitating the comparison of epigenetic profiles between gene pairs.

#### Feature Extraction using Random Projection for GSE76344 dataset

We investigate an alternative feature extraction approach utilizing random projection. The random projection is a method to reduce the dimensionality of the data by trading a controlled amount of accuracy (as additional variance) for faster processing times and smaller model size. The dimensions and distribution of random projections matrices are controlled so as to preserve the pairwise distances between any two samples of the dataset. Thus random projection is a suitable approximation technique for distance based method (14).We choose the Gaussian Random matrix projection for feature extraction. This will reduces the dimensionality by projecting the original input space on a randomly generated matrix where components are drawn from the following dis-tribution 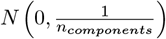.

#### Feature Extraction using Transformer

We utilize the `nn.TransformerEncoder` from Py-Torch (15). We define the parameter as follows: `d_model = 512, nhead = 8, num_layer = 6, output_dim = 20,000`. Where:

- `d_model:` the dimensionality of the model’s embeddings
- `nhead:` the number of independent attention mechanism within the multi-head attention layer
- `num_layer:` the number of sub-encoder-layers in the encoder
- `output_dim:` the output dimension

#### Feature Extraction using Auto Encoder (fully connected layer)

We build the autoencoder to flattened the input data into 1 × 20,000 dimensional vectors. The architecture is described in Table 2, where the `input_dim`is the input dimension, which is 20,000, and `bottleneck_dim`is the bottleneck dimension.

**Table 2.** Autoencoder Architecture using Fully Connected Layer

| Component | Layer | Linear Dimension | Batch Norm | Activation |
| --- | --- | --- | --- | --- |
| **Encoder** | Layer 1 | (input_dim, 1024) | 1024 | ReLU |
|  | Layer 2 | (1024, 512) | 512 | ReLU |
|  | Layer 3 | (512, 256) | 256 | ReLU |
|  | Layer 4 | (256, bottleneck_dim) | - | - |
| **Decoder** | Layer 1 | (bottleneck_dim, 256) | 256 | ReLU |
|  | Layer 2 | (256, 512) | 512 | ReLU |
|  | Layer 3 | (512, 1024) | 1024 | ReLU |
|  | Layer 4 | (1024, input_dim) | - | - |

#### Feature Extraction using Auto Encoder (convolution layer)

In the standard fully connected autoencoder implementation, input data is flattened into 1 × 20,000 dimensional vectors, which inevitably results in the loss of spatial information. To address this limitation, we implemented a convolutional autoencoder architecture that preserves the spatial relationships within the data. We used the 5 × 4, 000 matrix that represents five histone marks and 4,000 location (+/-2,000 bp from TSS). The network architecture is described in Table 3.

**Table 3.** Autoencoder Architecture using Convolutional Neural Network

| Component | Layer | Layer Type | Parameters | Batch Norm | Activation |
| --- | --- | --- | --- | --- | --- |
| **Encoder** | Layer 1 | Conv2D | (1, 16), kernel=(5,10), stride=(1,4), padding=(0,3) | 16 feature maps | ReLU |
|  | Layer 2 | Conv2D | (16, 32), kernel=(1,10), stride=(1,4), padding=(0,3) | 32 feature maps | ReLU |
|  | Layer 3 | Conv2D | (32, 64), kernel=(1,10), stride=(1,4), padding=(0,3) | 64 feature maps | ReLU |
|  | Layer 4 | ConvTranspose2D | (64, latent_dim), kernel=(1,5), stride=(1,2), padding=(0,2) | - | - |
| **Decoder** | Layer 1 | ConvTranspose2D | (latent_dim, 64), kernel=(1,5), stride=(1,2), padding=(0,2), output_padding=(0,1) | 64 feature maps | ReLU |
|  | Layer 2 | ConvTranspose2D | (64, 32), kernel=(1,10), stride=(1,4), padding=(0,3), output_padding=(0,2) | 32 feature maps | ReLU |
|  | Layer 3 | ConvTranspose2D | (32, 16), kernel=(1,10), stride=(1,4), padding=(0,3), output_padding=(0,0) | 16 feature maps | ReLU |
|  | Layer 4 | ConvTranspose2D | (16, 1), kernel=(5,10), stride=(1,4), padding=(0,3), output_padding=(0,0) | - | Sigmoid |

### Histone Marks Choice

We investigated the impact of each histone mark during the classification and pairwise ranking processes. It is hypothesized that various histone mark types exert distinct influences on gene expression levels, either repressing or promoting transcription. Ablation testing was employed to identify the histone marks that significantly contribute to determining gene expression levels. The ablation test was conducted in the GSE76344 datasets for pairwise ranking tasks.

### Pairwise Ranking

We investigated the potential for predicting relative gene expression levels based solely on histone modification markers. This study focused on developing a ranking model that could determine which genes that have higher expression levels by analyzing their associated histone modification patterns. Our experimental framework utilized the REMC E066 and GSE76344 datasets.

We partitioned the datasets into training (80%), validation (10%), and testing (10%) subsets. We implemented a strict non-leakage protocol wherein individual genes were confined to a single partition, thereby preventing any gene from appearing across multiple partitions and eliminating potential data contamination.

During the training, we randomly selected gene pairs from the training partition. For each pair, we assigned a binary classification label: ‘1’ if the expression level of the first gene exceeded that of the second gene, and ‘0’ otherwise. Model performance was subsequently evaluated on the validation partition, with final assessments conducted on the held-out testing partition to ensure generalization of our findings.

## Results

### Pairwise Ranking

The results of pairwise ranking task is shown in Table 4. For HepG2 GSE76344, the performance metrics reveal minimal differences among feature extraction methods (using full 20,000 features, random projection, Transformer, and both types of Auto Encoders), with test accuracies hovering around 70-71%. However, ranking with median value substantially improves performance (75.04% test accuracy), while the binary classification approach yields the best results for this dataset (76.82%).

**Table 4.** Comparison of different preprocessing techniques and pairwise ranking results across datasets

| Dataset | Preprocessing Technique | Validation accuracy (%) | Validation AUC | Test accuracy (%) | Test AUC |
| --- | --- | --- | --- | --- | --- |
| HepG2 (GSE76344) | Original features from +/- 2000 bp from TSS and five histones (20,000 features) | 70.82 | 0.71 | 71.46 | 0.71 |
|  | Feature extraction (random projection) | 70.12 | 0.70 | 70.61 | 0.71 |
|  | Feature extraction with Transformer | 70.67 | 0.70 | 71.05 | 0.71 |
|  | Feature extraction with Auto Encoder (fully connected layer) | 70.42 | 0.70 | 70.61 | 0.70 |
|  | Feature extraction with Auto Encoder (convolution layer) | 70.01 | 0.70 | 69.96 | 0.70 |
| REMC E066 | Windowing k = 100 of 10,000 bp (100 features) | 76.86 | 0.77 | 77.84 | 0.78 |

The REMC E066 dataset demonstrates superior performance overall, with its windowing approach (k=100 of 10,000 bp) achieving 77.84% test accuracy and its binary classification task reaching the highest metrics in the entire table (80.72% test accuracy, 0.81 test AUC). These results suggest that while complex neural architectures for feature reduction offer limited benefits, specialized ranking methods and task-specific approaches like binary classification significantly enhance prediction performance in genomic data analysis, with the REMC E066 dataset consistently outperforming HepG2 GSE76344 across comparable methods.

### Ablation Testing

The Table 5 shows ablation results for REMC E066 dataset with different item arrangements in a ranking task, displaying performance metrics across training, validation, and test sets. The REMC066 dataset demonstrates exceptional performance in histone mark-based ranking tasks, with the most complex marker combinations achieving test accuracies exceeding 77%. The top-performing combinations, such as H3K4me1-H3K4me3-H3K9me3-H3K27me3-H3K9ac-H3K27ac and H3K4me1-H3K4me3-H3K9me3-H3K27me3-H3K36me3-H3K9ac-H3K27ac, consistently deliver AUC values of 0.77, indicating strong discriminative power. Individual markers like H3K9me3 and H3K27me3 achieve lower results with test accuracies of 55.82% and 59.78% respectively, while mid-level combinations involving 3-4 markers typically reach the 74-75% accuracy range. The consistent alignment between training, validation, and test performance across all combinations indicates excellent model generalization, with minimal overfitting despite the complexity of some marker combinations.

**Table 5.**
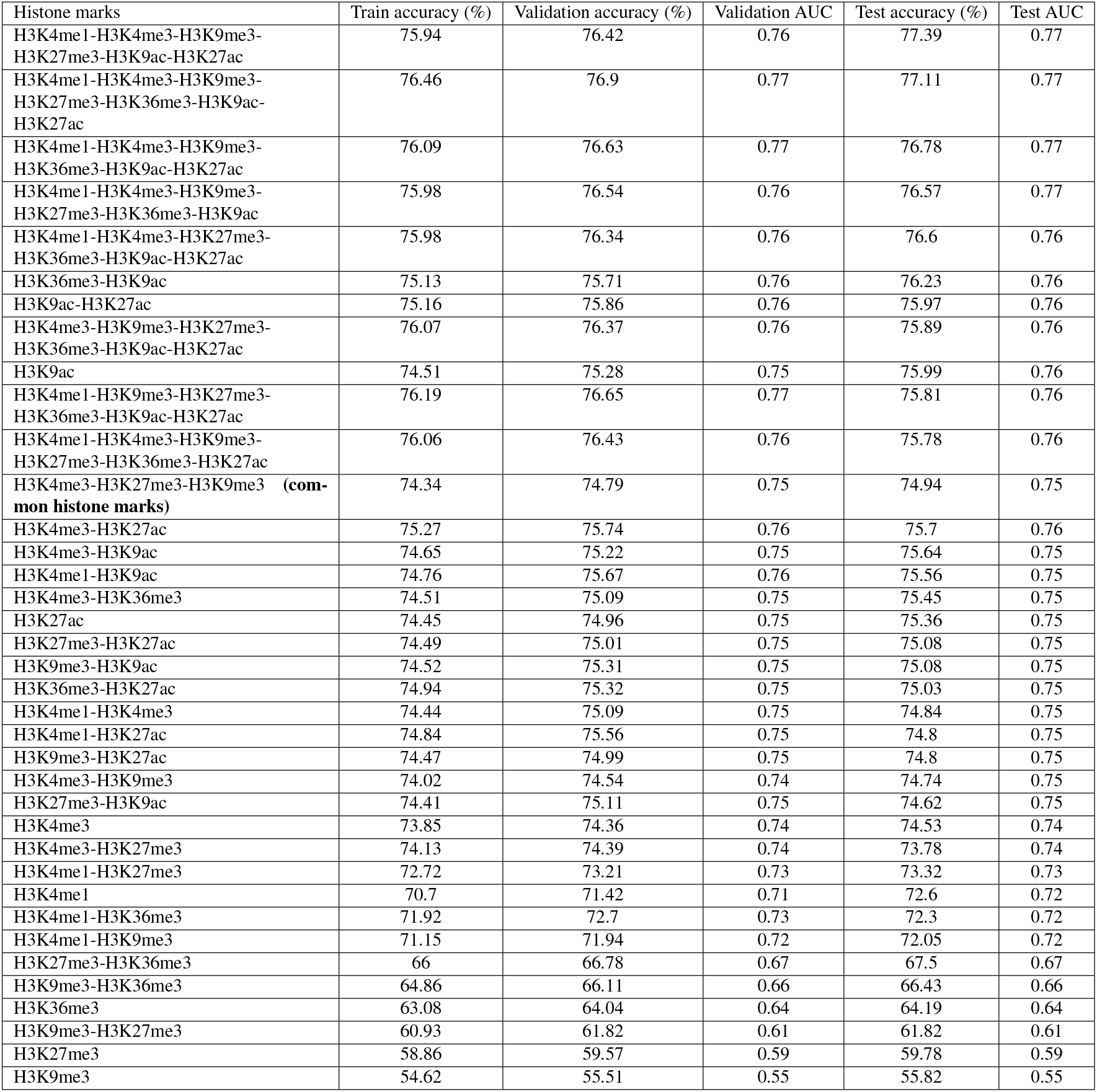
The Ablation Testing Result for Ranking using REMC E066 Dataset

| Histone marks | Train accuracy (%) | Validation accuracy (%) | Validation AUC | Test accuracy (%) | Test AUC |
| --- | --- | --- | --- | --- | --- |
| H3K4me1-H3K4me3-H3K9me3-H3K27me3-H3K9ac-H3K27ac | 75.94 | 76.42 | 0.76 | 77.39 | 0.77 |
| H3K4me1-H3K4me3-H3K9me3-H3K27me3-H3K36me3-H3K9ac-H3K27ac | 76.46 | 76.9 | 0.77 | 77.11 | 0.77 |
| H3K4me1-H3K4me3-H3K9me3-H3K36me3-H3K9ac-H3K27ac | 76.09 | 76.63 | 0.77 | 76.78 | 0.77 |
| H3K4me1-H3K4me3-H3K9me3-H3K27me3-H3K36me3-H3K9ac | 75.98 | 76.54 | 0.76 | 76.57 | 0.77 |
| H3K4me1-H3K4me3-H3K27me3-H3K36me3-H3K9ac-H3K27ac | 75.98 | 76.34 | 0.76 | 76.6 | 0.76 |
| H3K36me3-H3K9ac | 75.13 | 75.71 | 0.76 | 76.23 | 0.76 |
| H3K9ac-H3K27ac | 75.16 | 75.86 | 0.76 | 75.97 | 0.76 |
| H3K4me3-H3K9me3-H3K27me3-H3K36me3-H3K9ac-H3K27ac | 76.07 | 76.37 | 0.76 | 75.89 | 0.76 |
| H3K9ac | 74.51 | 75.28 | 0.75 | 75.99 | 0.76 |
| H3K4me1-H3K9me3-H3K27me3-H3K36me3-H3K9ac-H3K27ac | 76.19 | 76.65 | 0.77 | 75.81 | 0.76 |
| H3K4me1-H3K4me3-H3K9me3-H3K27me3-H3K36me3-H3K27ac | 76.06 | 76.43 | 0.76 | 75.78 | 0.76 |
| H3K4me3-H3K27me3-H3K9me3 **(common histone marks)** | 74.34 | 74.79 | 0.75 | 74.94 | 0.75 |
| H3K4me3-H3K27ac | 75.27 | 75.74 | 0.76 | 75.7 | 0.76 |
| H3K4me3-H3K9ac | 74.65 | 75.22 | 0.75 | 75.64 | 0.75 |
| H3K4me1-H3K9ac | 74.76 | 75.67 | 0.76 | 75.56 | 0.75 |
| H3K4me3-H3K36me3 | 74.51 | 75.09 | 0.75 | 75.45 | 0.75 |
| H3K27ac | 74.45 | 74.96 | 0.75 | 75.36 | 0.75 |
| H3K27me3-H3K27ac | 74.49 | 75.01 | 0.75 | 75.08 | 0.75 |
| H3K9me3-H3K9ac | 74.52 | 75.31 | 0.75 | 75.08 | 0.75 |
| H3K36me3-H3K27ac | 74.94 | 75.32 | 0.75 | 75.03 | 0.75 |
| H3K4me1-H3K4me3 | 74.44 | 75.09 | 0.75 | 74.84 | 0.75 |
| H3K4me1-H3K27ac | 74.84 | 75.56 | 0.75 | 74.8 | 0.75 |
| H3K9me3-H3K27ac | 74.47 | 74.99 | 0.75 | 74.8 | 0.75 |
| H3K4me3-H3K9me3 | 74.02 | 74.54 | 0.74 | 74.74 | 0.75 |
| H3K27me3-H3K9ac | 74.41 | 75.11 | 0.75 | 74.62 | 0.75 |
| H3K4me3 | 73.85 | 74.36 | 0.74 | 74.53 | 0.74 |
| H3K4me3-H3K27me3 | 74.13 | 74.39 | 0.74 | 73.78 | 0.74 |
| H3K4me1-H3K27me3 | 72.72 | 73.21 | 0.73 | 73.32 | 0.73 |
| H3K4me1 | 70.7 | 71.42 | 0.71 | 72.6 | 0.72 |
| H3K4me1-H3K36me3 | 71.92 | 72.7 | 0.73 | 72.3 | 0.72 |
| H3K4me1-H3K9me3 | 71.15 | 71.94 | 0.72 | 72.05 | 0.72 |
| H3K27me3-H3K36me3 | 66 | 66.78 | 0.67 | 67.5 | 0.67 |
| H3K9me3-H3K36me3 | 64.86 | 66.11 | 0.66 | 66.43 | 0.66 |
| H3K36me3 | 63.08 | 64.04 | 0.64 | 64.19 | 0.64 |
| H3K9me3-H3K27me3 | 60.93 | 61.82 | 0.61 | 61.82 | 0.61 |
| H3K27me3 | 58.86 | 59.57 | 0.59 | 59.78 | 0.59 |
| H3K9me3 | 54.62 | 55.51 | 0.55 | 55.82 | 0.55 |

The Table 6 shows ablation results for GSE76344 dataset with different item arrangements in a ranking task, displaying performance metrics across training, validation, and test sets. The GSE76344 dataset presents more challenging ranking conditions compared to REMC066, with peak performance reaching approximately 71% test accuracy. The best-performing marker combinations, including H3K9me3-H3K27ac-H3K27me3-H3K4me3-H3K9ac and H3K4me3-H3K27me3-H3K9me3-H3K27ac, achieve test AUC values of 0.71. The ablation results reveal a steep performance decline as marker complexity decreases, with individual markers performing particularly poorly. Single markers like H3K9me3 and H3K27me3 show substantially reduced effectiveness with test accuracies of only 53.43% and 55.11% respectively, barely exceeding random classification performance.

**Table 6.** The Ablation Testing Result for Ranking using GSE76344 Dataset

| Histone marks | Train accuracy (%) | Validation accuracy (%) | Validation AUC | Test accuracy (%) | Test AUC |
| --- | --- | --- | --- | --- | --- |
| H3K9me3-H3K27ac-H3K27me3-H3K4me3-H3K9ac | 71.2 | 70.44 | 0.7 | 71.72 | 0.71 |
| H3K4me3-H3K27me3-H3K9me3-H3K27ac | 71.12 | 70.54 | 0.7 | 71.29 | 0.71 |
| H3K27me3-H3K9ac-H3K4me3-H3K27ac | 71.26 | 70.46 | 0.7 | 71.25 | 0.71 |
| H3K9ac-H3K27me3-H3K9me3-H3K27ac | 70.55 | 70.17 | 0.7 | 71.01 | 0.71 |
| H3K9me3-H3K4me3-H3K27ac-H3K9ac | 70.86 | 70.42 | 0.7 | 71.1 | 0.71 |
| H3K27me3-H3K9ac-H3K4me3-H3K9me3 | 70.6 | 69.93 | 0.69 | 71.04 | 0.71 |
| H3K4me3-H3K27me3-H3K9me3 (**common histone marks**) | 70.55 | 70.03 | 0.70 | 69.61 | 0.69 |
| H3K9me3-H3K9ac | 69.94 | 69.84 | 0.69 | 70.42 | 0.7 |
| H3K27ac-H3K4me3 | 70.88 | 70.23 | 0.7 | 70.45 | 0.7 |
| H3K9ac-H3K27ac | 70.42 | 70.19 | 0.7 | 70.17 | 0.7 |
| H3K9me3-H3K4me3 | 69.89 | 69.54 | 0.69 | 70.13 | 0.7 |
| H3K9ac-H3K4me3 | 70.59 | 70 | 0.7 | 69.82 | 0.69 |
| H3K4me3-H3K27me3 | 70.04 | 69.48 | 0.69 | 69.77 | 0.69 |
| H3K27me3-H3K27ac | 69.65 | 69.49 | 0.69 | 69.85 | 0.69 |
| H3K27me3-H3K9ac | 70.03 | 69.86 | 0.69 | 69.38 | 0.69 |
| H3K9me3-H3K27ac | 69.66 | 69.56 | 0.69 | 69.44 | 0.69 |
| H3K9me3-H3K27me3 | 55.11 | 55.05 | 0.53 | 55.36 | 0.53 |
| H3K9ac | 69.88 | 69.62 | 0.69 | 69.68 | 0.69 |
| H3K4me3 | 70 | 69.4 | 0.69 | 69.17 | 0.69 |
| H3K27ac | 69.6 | 69.49 | 0.69 | 69.34 | 0.69 |
| H3K27me3 | 54.88 | 54.9 | 0.53 | 55.11 | 0.53 |
| H3K9me3 | 52.7 | 52.85 | 0.5 | 53.43 | 0.5 |

## Conclusions

Our research demonstrates successful processing of the GSE76344 dataset, making it viable for diverse machine learning applications in genomic analysis. The histone mark data proved effective in determining gene expression levels. The pairwise ranking methodology showed promising performance, offering an alternative analytical framework that may be advantageous in certain contexts. Importantly, our models showed promising signs of generalization capabilities, indicating their potential utility beyond the specific datasets examined. These findings advance our understanding of how epigenetic markers can be effectively leveraged to predict gene expression states and demonstrate computational approaches for analyzing histone modification data.

Our findings highlight several avenues for future research. First, expanding into cross-cell analysis represents a critical research direction; our current study is limited to HepG2 GSE76344 and REMC E066, which constrains the generalizability of our conclusions. Second, integration with additional epigenetic factors, such as DNA methylation and non-coding RNA, could provide a more comprehensive understanding of gene regulation mechanisms. Such a multi-omics approach would likely yield more robust predictive models by capturing the complex interplay between various epigenetic modifications that collectively influence gene expression patterns. These proposed extensions would address the current limitations of our work while advancing the field’s understanding of epigenetic regulation across different cellular contexts.

## ACKNOWLEDGEMENTS

Acknowledgement

